# A Quantitative Review of Biodiversity Trends during the Great Ordovician Biodiversification Event

**DOI:** 10.1101/2025.11.05.686803

**Authors:** Jared Cliff Richards, Karma Nanglu, Javier Ortega-Hernández

**Author notes:** **Author Contributions:** Jared Richards conducted the literature search and data analysis. Jared Richards, Karma Nanglu, and Javier Ortega-Hernández contributed to the interpretation of results, discussion, and writing. All authors approved the final manuscript. **Competing Interest Statement:** There are no competing interests.

## Abstract

The Great Ordovician Biodiversification Event (GOBE) embodies the most dramatic increase of marine biodiversity and escalation of macroecological complexity during the early Phanerozoic. Despite its critical role, the precise timing and duration of the GOBE remain controversial. Numerous palaeobiological studies have attempted to quantify the GOBE based on estimates of species richness through time for various groups of marine organisms. However, these studies use fossil data restricted to specific geographic regions or employ disparate methodologies that preclude direct analytical comparisons. We present a meta-analysis of Ordovician biodiversity that integrates information from multiple temporal, geographic, and ecological scales. We collate 98 datasets from 54 publications to analyze temporally standardized rates of marine species biodiversity accumulation between the latest Cambrian and throughout the entire Ordovician using an effect-size approach. Our results indicate statistically significant high rates of sustained species accumulation that can be traced from the late Cambrian and until the Middle Ordovician, stabilization during the Late Ordovician and then a precipitous decline caused by the Late Ordovician Mass Extinction. Geographic scale (global vs regional) has no significant bearing on rates of biodiversification, with the only exception observed during the Dapingian-Darriwilian transition, supporting the hypothesis of mass dispersal of generalists during the Early Ordovician. Benthic and suspension-feeding organisms show high rates of biodiversity accumulation throughout most of the Ordovician (Tremadocian-Sandbian), whereas the diversification of nektonic, pelagic and predatory/scavenger organisms was mostly restricted to the Early Ordovician.

## Introduction

The Great Ordovician Biodiversification Event, or GOBE, comprises a series of major evolutionary radiations at low taxonomic levels across multiple marine animal clades paired with substantial increases in ecological complexity due to increased suspension feeding and ecological tiering, leading to more intricate food webs (1, 2). Despite its critical role in the emergence of modern marine biodiversity and ecosystem structure, the precise timing and duration of the GOBE remains elusive. Although numerous paleobiological studies that quantify biodiversity through time have explored the onset and duration of the GOBE (1–4) (Fig. 1A), they vary drastically in their geographic range, geographic scale, clades included, and diversity metrics. The breadth of these studies has not produced a consensus regarding the onset or duration of the GOBE. It has been suggested that the GOBE was restricted to the Middle Ordovician based on an observed maximum species richness during the Darriwilian (3–5) (Fig. 1A). Alternatively, the GOBE could be a more protracted evolutionary event that began during the Cambrian consisting of a series of diversifications throughout the Ordovician that asynchronously affected different clades and ecological guilds (1, 3–7). This matter is further complicated by the use of disparate methodologies that preclude direct comparisons between the results of different studies (8) and hinder a consolidated perspective on this major evolutionary radiation. Here, we employ an effect-size approach to explore the nature of the GOBE, which allows us to statistically compare rates of biodiversity change through deep time. By standardizing rates of biodiversity accumulation between consecutive geological stages and series, we synthesize information from multiple datasets into a single framework that allows us to illuminate broad patterns of biodiversity changed from the latest Cambrian and the entire Ordovician.

**Figure 1.**
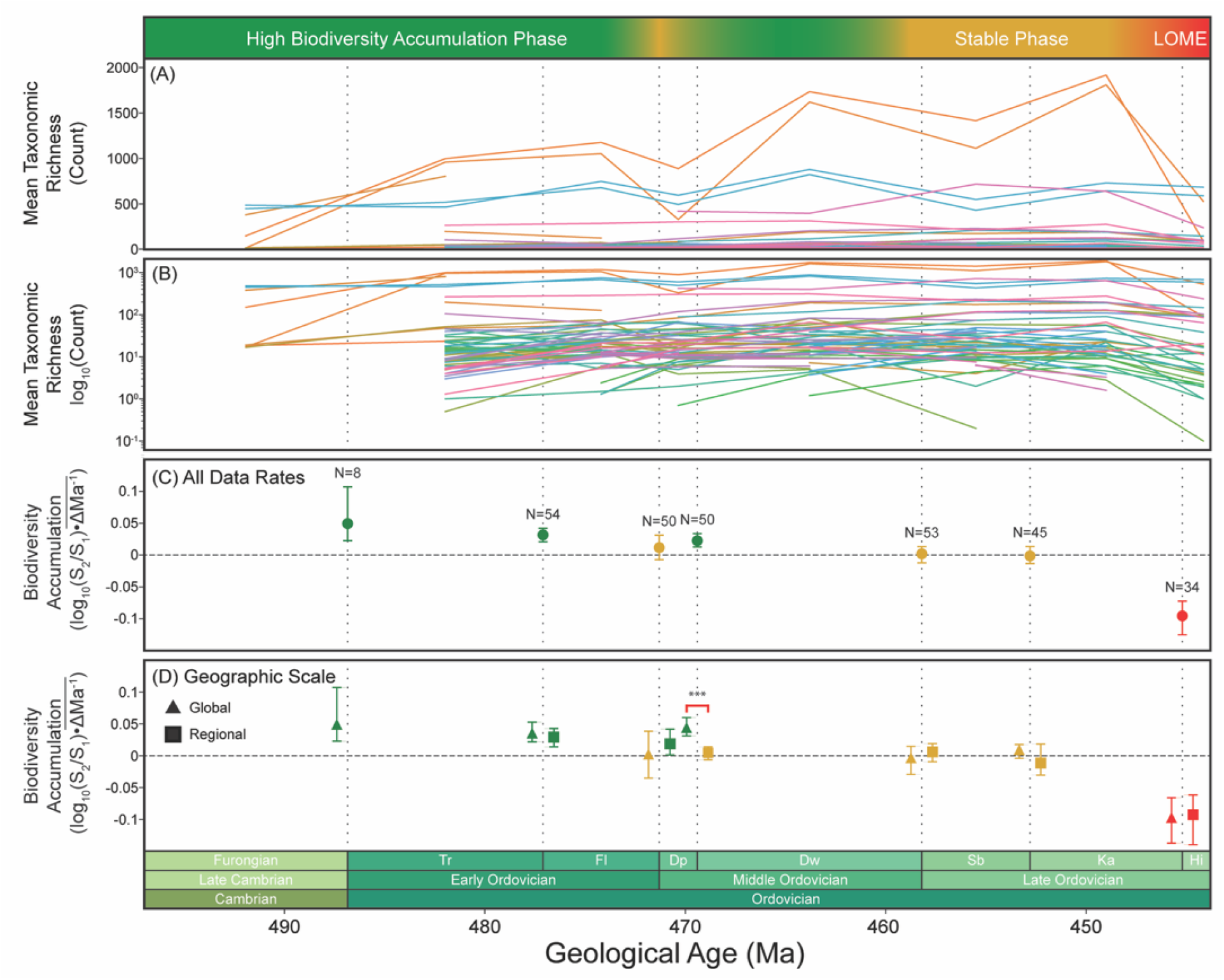
General data and methodological assessments of biodiversification rates. A) Reconstructed diversity curves used for meta-analysis showing raw taxonomic counts. B) Reconstructed diversity curves used for meta-analysis depicted on a logarithmic scale. C) Standardized rate of biodiversity accumulation. D) Standardized rate of biodiversity accumulation separated by geographic scale (regional vs. global). Symbol coloring indicates positive (green), neutral (yellow), or negative (red) rates of biodiversity change at a given transition.

## Methods

We searched for peer-reviewed studies recording early Paleozoic diversity using the Web of Science on August 15^th^, 2024, using the following criteria: [TITLE-ABS-KEY (“Cambrian” OR “Ordovician”) AND biodiversi* OR diversi* OR richness OR composition OR communit*)]. These search parameters, which cover all studies reporting diversity data throughout the Cambrian and Ordovician, produced 8,634 results, which were further vetted by retaining articles that provide information on richness or biodiversity throughout discrete units of geologically relevant time, and discarding all others. Given the paucity of publicly available biodiversity data for the Cambrian that met the above criteria, we restricted our analysis from the latest Cambrian to the end-Ordovician. These criteria resulted in a final total of 54 research papers encompassing 98 datasets being included in all subsequent analyses.

The resultant studies from the above search were examined for documented biodiversity values spanning consecutive geologic timeframes, specifically stages or series, to calculate the effect size of diversification rates between contiguous temporal units. We used PlotDigitizer 3.3.9 (https://plotdigitizer.com/) to extract and record the relevant biodiversity information from each compatible paper alongside the diversity metric used, taxonomic clade, taxonomic level of recorded diversity, ecological tiering, feeding strategy, and geographic scale, when available.

For each dataset and temporal transition based on count data, the temporally standardized rate of biodiversity accumulation between successive time frames was calculated as: 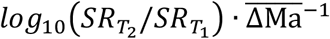 Where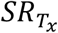 is the biodiversity at a given temporal interval, and 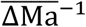 is the average duration of the two consecutive time frames in millions of years. This metric, refined from modern macroecological research (9), provides a metric to examine the magnitude and direction of change from one time to the next for each study and allows us to standardize for the wide variance in the early Paleozoic stage and series duration. To analyze the full dataset and the data grouped by geographic scale, we calculated the 95% confidence interval of the mean via bias-corrected and accelerated bootstrapping (10,000 iterations). Given the smaller sample sizes, we used boxplots to show the location and spread of the temporally standardized rate of biodiversity accumulation grouped by ecological tiering and feeding strategy (10).

All data analysis, statistics, simulations, and visualizations were completed in Visual Studio Code using Python 3.12.0 and the pandas, random, math, numPy, matplotlib, statsmodel, seaborn, and sciPy.

## Results

We first reconstructed diversity curves for each study (Fig. 1A, B), recapitulating the conventional methods used to represent diversity through time (1–8, 11, 12) to facilitate a visual comparison with the results of the meta-analysis. To statistically analyze diversification patterns for the effect-size approach, we bootstrapped 95% CI for the mean biodiversity accumulation rates. The Furongian–Tremadocian, Tremadocian–Floian, and Dapingian–Darriwilian transitions are bounded above zero, signifying positive biodiversity accumulation (Fig. 1C). The bootstrapped 95% CI for the Floian–Dapingian, Dapingian– Sandbian, and Sandbian–Katian transitions overlap with zero, signifying that rates of biodiversity accumulation remained stable at these points (Fig. 1C). The bootstrapped 95% CI for the Katian–Hirnantian transition is bounded below zero, signifying a decrease in biodiversity (*i*.*e*., mass extinction) (Fig. 1B). We explored the effects of geographic scale by subdividing data into regional and global partitions across the same geological transitions (Fig. 1D). The results indicate minimal discrepancies between these geographic scales, with the only timeframe where global and regional rates of biodiversity accumulation differ statistically is during the Dapingian-Darriwilian transition, where global rates are significantly higher (Fig. 1D).

We then explored the diversification of major ecological guilds (Fig. 2, Supplementary Table 1). Biodiversification rates for benthic clades (e.g. trilobites, brachiopods, echinoderms) are positive and increase from the Tremadocian until the Sandbian–Katian transition, where they stagnate and eventually become negative during the Katian–Hirnantian transition (Fig. 2A). Rates for nektonic and planktonic clades (e.g. cephalopods, conodonts, phyto- and zooplankton) follow each other throughout the Ordovician, with a positive trend during the Tremadocian–Floian and Dapingian–Darriwilian transitions, but negative during all other transitions from the Middle to Late Ordovician (Fig. 2A). Suspension feeders have positive rates throughout the Tremadocian until Darriwilian (Early and Middle Ordovician); these rates stagnate and become negative during the Sandbian–Katian and Katian–Hirnantian (Late Ordovician), respectively (Fig. 2B). Predators and scavengers only exhibit positive rates of biodiversity accumulation during the Tremadocian– Floian (Early Ordovician), whereas their rates stagnated during the Dapingian–Darriwilian and were negative during all subsequent transitions (Fig. 2B).

**Figure 2.**
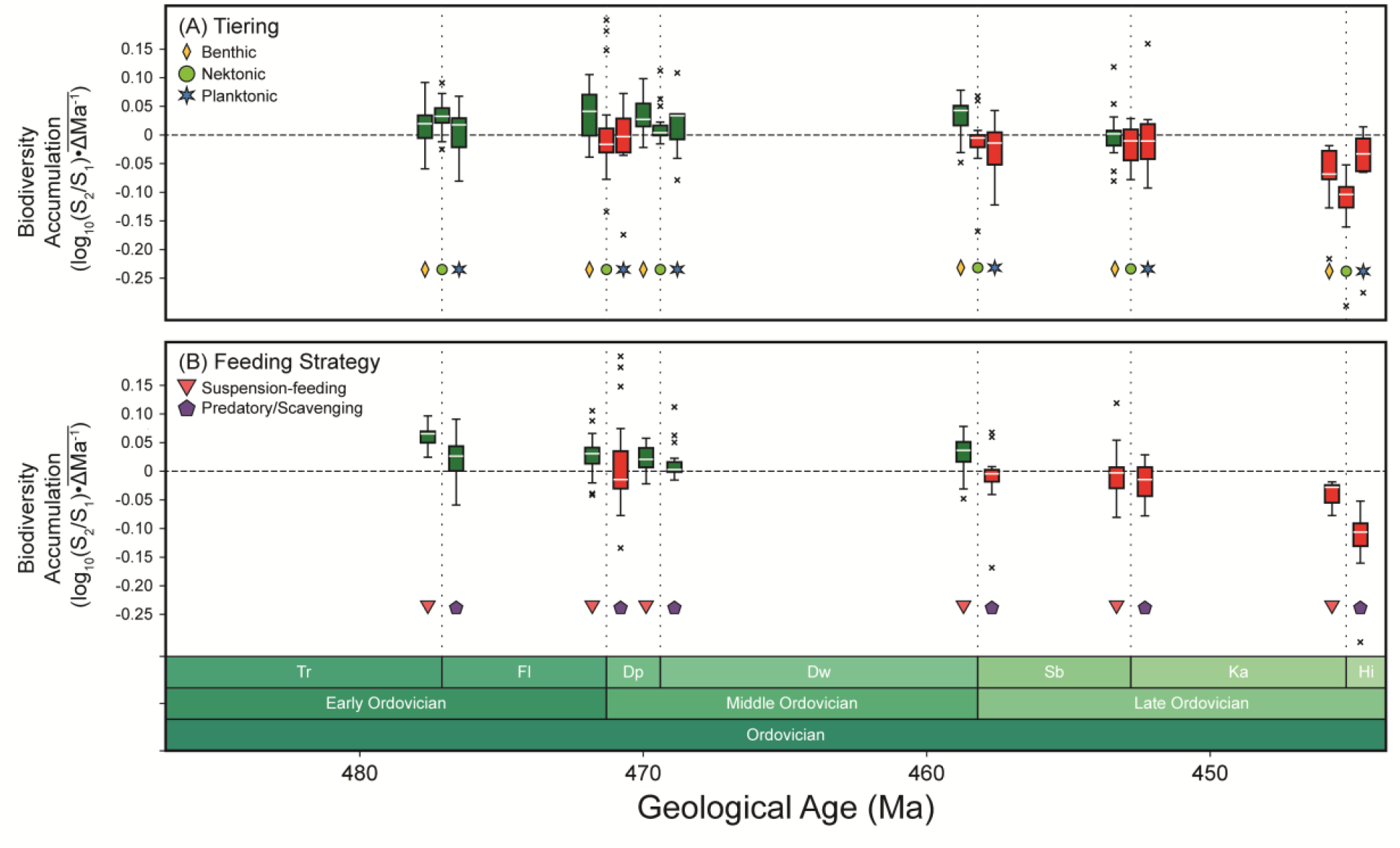
Ecological assessment of biodiversification rates. A) Standardized rate of biodiversity accumulation separated by ecological tiering. B) Standardized rate of biodiversity accumulation separated by feeding strategy. Symbol coloring indicates positive (green), or negative (red) rates of biodiversity change at a given timeframe.

## Discussion

The GOBE has been considered as the climax to a series of background early Ordovician Radiations from the Tremadocian to Dapingian defined by slow biodiversity accumulation, with the GOBE proper taking place from the Dapingian-Darriwilian until the early Katian and ending with the Late Ordovician Mass Extinction (LOME) during the late Katian and Hirnatian stages (1, 3, 6) (Fig. 1A). Our findings directly challenge this notion by showing that the Early and Middle Ordovician, specifically from the Furongian-Tremadocian to the Dapingian-Darriwilian transitions, experienced the highest rates of biodiversity accumulation for the entire Ordovician (1, 3, 5) (Fig. 1C), with the exception of the Floian-Dapingian boundary. Although the Dapingian-Darriwilian transition is commonly regarded as the start of the main pulse of the GOBE based on taxonomic counts (6), this timepoint did not show uniquely higher rates of biodiversity accumulation relative to the Early Ordovician. Instead, the sharp increase in marine species observed in taxonomic counts is likely a consequence of the cumulative effect of the high rates of biodiversity accumulation during the Early Ordovician. The Darriwilian-Sandbian and Sandbian-Katian transitions signal intervals of neutral biodiversity. When focusing on the rates of biodiversity accummulation, rather than raw taxonomic counts, it is clear that the dramatic radiation that typifies the GOBE was effectively active for the entire Early Ordovician, rather than just starting in the Middle Ordovician (3, 4, 6) (Fig. 1C). Critically, our results bridge the purported “biodiversity gap” between the late Cambrian and the Middle Ordovician and support the GOBE as a long-term and large-scale Paleozoic radiation (1, 3). The substantial rate of biodiversity accumulation that typifies the GOBE did not culminate in the Katian, contrary to the pattern observed using raw taxonomic counts alone (6) (Fig. 1A). Instead, we find that biodiversity accumulation rates had already become neutral during the Darriwilian-Sandbian and remained stable during the Sandbian-Katian (Fig. 1C), followed by the catastrophic negative rate of biodiversity accumulation during the Katian-Hirnatian caused from by the LOME (13).

Our results also allow us to examine how geographic scale affects our understanding of the GOBE. We find that global and regional-scale rates of biodiversity accumulation were statistically indistinguishable in most transitions, with the Dapingian-Darriwilian as the only exception (Fig. 1D). Global rates of biodiversity accumulation consistently exceeded regional rates in terms of magnitude (Fig. 1D), which concurs with previous findings that the GOBE results from a series of asynchronous radiations at local and regional scales, that when combined, manifest as dramatic global-scale fluctuations in net biodiversity throughout the Ordovician (1, 4, 5, 8). The Dapingian-Darriwilian transition shows a significant discrepancy between a positive rate of biodiversity accumulation at the global scale, but neutral rate of accumulation at the regional scale (Fig. 1D). In this context, the Dapingian-Darriwilian correlates with the drastic increase of dispersal of ecologically generalist organisms that would outcompete and accelerate the extirpation of local specialist taxa (8, 14, 15). This discrepancy highlights the fact that despite the significant deceleration of rates of biodiversity accumulation at regional levels during the Dapingian-Darriwilian, likely owing to the increased ecological competition due to higher dispersal at the time (6), the aggregate global signal still maintained a positive rate of biodiversity accumulation overall.

The sequence of diversifications of primarily planktonic (late-Cambrian to Early Ordovician) and benthic communities (Early to Middle Ordovician) are a key component of the GOBE’s paleoecology (1, 3, 5). Our results broadly capture some of these diachronous patterns but refine their timing (Fig. 2). All ecological guilds showed positive rates of biodiversity accumulation during the Tremadocian-Floian, but only benthic organisms continuously radiated into the Late Ordovician (Sandbian-Katian) (Fig. 2A). By contrast, nekton (*e*.*g*., cephalopods, conodonts) and plankton (*e*.*g*., acritarchs, graptolites) closely mirrored each other undergoing negative rates of biodiversity accumulation during the Floian-Dapingian, recovery during the apingian-Darriwilian, and consistent loss for the remainder of the Middle-Late Ordovician. Suspension feeding organisms maintained positive rates of biodiversity accumulation until the Middle Ordovician and only decreased during the Sandbian-Katian and subsequently the LOME, but predators/scavengers fluctuated during the Middle Ordovician and then experienced losses until the end of the Late Ordovician (Fig. 2B). Taken together, these patterns provide quantitative support for a fundamental ecological decoupling between the biodiversification dynamics of seafloor and the water column communities that came fully consolidated during the Middle Ordovician.

In conclusion, we posit that traditional taxonomic counts do not accurately capture the nuance of the GOBE as a large-scale and long-term radiation because of their emphasis on marine species diversity (Fig. 1A), rather than the speed of the radiation itself. By focusing on standardized rates of biodiversity accumulation (Fig. 1C) we can mitigate the effect-size bias caused by raw taxonomic counts. Our findings provide evidence that the interval between the Furongian to Dapingian embody the fastest diversification episode of the GOBE, but this effect was understated by the relatively low species richness at the time, particularly when compared to the peak in with global biodiversity reached during the Middle and Late Ordovician.

## Notes

### Competing Interest Statement

The authors have declared no competing interest.

### Funding

- NSF: 2047192, 2140743

## References

1. T. Servais, D. A. T. Harper, The Great Ordovician Biodiversification Event (GOBE): definition, concept and duration. LET 51, 151–164 (2018).

2. J. Fan, et al., A high-resolution summary of Cambrian to Early Triassic marine invertebrate biodiversity. Science 367, 272–277 (2020).

3. T. Servais, et al., No (Cambrian) explosion and no (Ordovician) event: A single long-term radiation in the early Palaeozoic. Palaeogeography, Palaeoclimatology, Palaeoecology 623, 111592 (2023).

4. A. L. Stigall, C. T. Edwards, R. L. Freeman, C. M. Ø. Rasmussen, Coordinated biotic and abiotic change during the Great Ordovician Biodiversification Event: Darriwilian assembly of early Paleozoic building blocks. Palaeogeography, Palaeoclimatology, Palaeoecology 530, 249–270 (2019).

5. T. Servais, A. W. Owen, D. A. T. Harper, B. Kröger, A. Munnecke, The Great Ordovician Biodiversification Event (GOBE): The palaeoecological dimension. Palaeogeography, Palaeoclimatology, Palaeoecology 294, 99–119 (2010).

6. A. L. Stigall, R. L. Freeman, C. T. Edwards, C. M. Ø. Rasmussen, A multidisciplinary perspective on the Great Ordovician Biodiversification Event and the development of the early Paleozoic world. Palaeogeography, Palaeoclimatology, Palaeoecology 543, 109521 (2020).

7. Y. Deng, et al., Timing and patterns of the Great Ordovician Biodiversification Event and Late Ordovician mass extinction: Perspectives from South China. Earth-Science Reviews 220, 103743 (2021).

8. F. Saleh, J. B. Antcliffe, L. Lustri, A. C. Daley, C. Gibert, Cambrian and Ordovician diversity fluctuations could be resolved through a single ecological hypothesis. LET 56, 1–13 (2023).

9. M. Vellend, et al., Global meta-analysis reveals no net change in local-scale plant biodiversity over time. Proc. Natl. Acad. Sci. U.S.A. 110, 19456–19459 (2013).

10. M. Puth, M. Neuhäuser, G. D. Ruxton, On the variety of methods for calculating confidence intervals by bootstrapping. Journal of Animal Ecology 84, 892–897 (2015).

11. C. T. Edwards, Links between early Paleozoic oxygenation and the Great Ordovician Biodiversification Event (GOBE): A review. Palaeoworld 28, 37–50 (2019).

12. A. L. Stigall, Ordovician oxygen and biodiversity. Nature Geosci 10, 887–888 (2017).

13. D. P. G. Bond, S. E. Grasby, Late Ordovician mass extinction caused by volcanism, warming, and anoxia, not cooling and glaciation. Geology 48, 777–781 (2020).

14. T. Servais, T. Danelian, D. A. T. Harper, A. Munnecke, Possible oceanic circulation patterns, surface water currents and upwelling zones in the Early Palaeozoic. GFF 136, 229–233 (2014).

15. F. Saleh, et al., Contrasting Early Ordovician assembly patterns highlight the complex initial stages of the Ordovician Radiation. Sci Rep 12, 3852 (2022).

